# Exploring the Biochemical Landscape of Bacterial Medium with Pyruvate as the Exclusive Carbon Source for NMR Studies

**DOI:** 10.1101/2024.10.26.620443

**Authors:** Çağdaş Dağ, Kerem Kahraman

## Abstract

The use of *Escherichia coli* for recombinant protein production is a cornerstone in structural biology, particularly for nuclear magnetic resonance (NMR) spectroscopy studies. Understanding the metabolic behavior of *E. coli* under different carbon sources is critical for optimizing isotope labeling strategies, which are essential for protein structure determination by NMR. Recent advancements, such as mixed pyruvate labeling, have enabled improved backbone resonance assignment in large proteins, making selective isotopic labeling strategies more important than ever for NMR studies. In this study, we aimed to investigate the metabolic adaptations of *E. coli* when grown on pyruvate as the sole carbon source, a common condition used to achieve selective labeling for NMR spectroscopy. Using NMR-based metabolomics, we tracked key metabolic shifts throughout the culture process to better understand how pyruvate metabolism affects protein production and isotopic labeling. Our results reveal that pyruvate is rapidly depleted before IPTG induction, while acetate and lactate accumulate due to overflow metabolism. These byproducts persist after induction, indicating that pyruvate is diverted into waste pathways, which limits its efficient use in isotope incorporation. This metabolic inefficiency presents a challenge for isotopic labeling protocols that rely on pyruvate as a carbon source for NMR studies. Our results highlight the need to fine-tune pyruvate supplementation to improve metabolic efficiency and isotopic labeling, making this study directly relevant to optimizing protocols for NMR studies involving protein structure determination. These insights provide valuable guidance for enhancing the quality and yield of isotopically labeled proteins in NMR spectroscopy.

## Introduction

Recombinant protein production is a crucial technique for conducting research on proteins. The desired protein can be produced in a laboratory setting using organisms such as *Escherichia coli* and, after purification, can be used in various studies. The DNA containing the gene encoding the protein is first isolated from the relevant organism and then cleaved by restriction enzymes. Similarly, the plasmid to be used as a vector is also isolated and cleaved using the same restriction enzymes (Nathans & Smith, 1975). This ensures compatibility between the cut sites of the vector and the DNA to be inserted. The ends of the DNA and the vector are then ligated using DNA ligase enzyme. As a result of this process, the recombinant plasmids are prepared for use and transferred into bacteria capable of transformation (Nathans & Smith, 1975). Successfully transformed bacteria are grown in appropriate culture media, and protein production is induced once a sufficient number of bacteria have been obtained. After a certain period, the cells present in the culture are harvested by centrifugation, lysed, and the produced protein is purified.

One of the methods used to determine the structures of the produced and purified proteins is nuclear magnetic resonance (NMR) spectroscopy (Wüthrich, 1990). In NMR spectroscopy, a strong magnetic field is applied to the sample in solution. As a result, the nuclei of the atoms align with the magnetic field. The nuclei can align either in the same or opposite direction to the magnetic field, placing each nucleus in either a high-energy or low-energy state. Various radiofrequency signals are then transmitted to excite the nuclei in low-energy states to higher energy levels. During this transition, each nucleus absorbs radio waves with specific energy and later emits this energy as radio waves when returning to the low-energy state (Gerothanassis et al., 2002). The energy absorbed by the nuclei varies depending on the strength of the magnetic field applied, the type of atom being excited, and the atoms to which the nucleus is bonded or interacting with (Cavalli et al., 2007). The frequencies at which the sample absorbs energy are detected, and the chemical shifts relative to a reference compound are calculated (Cavalli et al., 2007). These calculated chemical shifts are represented on a graph, and peaks are observed at the frequencies where NMR signals are obtained. By examining the positions of these peaks, the structure of small molecules can be determined.

Since not all atomic nuclei are NMR-active, it is necessary to use specific isotopes in the sample. For a nucleus to be NMR-active, its spin must be greater than zero. Although some nuclei with integer spins are NMR-active, nuclei with half-integer spins, such as ^1^H, ^13^C, and ^15^N, produce the most useful signals due to their narrow absorption ranges (Gerothanassis et al., 2002). ^1^H is the most common isotope of hydrogen and is found in all organic compounds, making hydrogen NMR one of the most widely used techniques. In small molecules with few atoms, data collected using only hydrogen atoms is sufficient for structural determination. As long as the peaks are sufficiently separated, a structural model can be constructed based on the interactions between hydrogen atoms. However, as the size of the molecule increases, such as in proteins, the number of atoms also increases, making measurements using only hydrogen insufficient for structural determination. Since proteins are composed of thousands of atoms, analyzing such complex structures by relying solely on hydrogen atom signals is not feasible (Wüthrich, 1990). Therefore, the use of other NMR-active nuclei is essential. The most abundant elements in proteins are carbon, nitrogen, hydrogen, and oxygen. However, the most common carbon isotope, ^12^C, does not have an NMR-active nucleus. Similarly, ^14^N, the most common nitrogen isotope, is not useful for protein structural determination due to its wide NMR signal range. As a result, isotopically labeled proteins must be produced to gather detailed information about a protein using NMR (Wüthrich, 1990).

Isotopically labeled protein production follows a different path than standard recombinant protein production protocols. In recombinant protein production, nutrient-rich media are used to ensure that bacteria proliferate rapidly and produce large amounts of protein. These nutrient-rich media typically contain amino acids, salts, carbohydrates as a carbon and energy source, vitamins, metal ions, and nucleic acids. Bacteria can directly use the amino acids present in the medium for protein production. However, this is undesirable for the production of isotopically labeled proteins. To address this, minimal media are employed in isotopic labeling processes. Unlike rich media, minimal media do not contain amino acids. Instead, compounds containing elements such as nitrogen, carbon, sulfur, calcium, and sodium are added along with a carbon source (e.g., glucose) (Azatian et al., 2019). This approach forces the bacteria to synthesize their own amino acids. Rather than adding pre-labeled amino acids to the medium, this method is more practical. Any of the minimal medium components can be added as isotopically labeled, allowing for the production of proteins with specific labeled atoms. For example, labeled NH_4_Cl is added for ^15^N-labeled proteins, while labeled glucose is used for ^13^C-labeled proteins (Cai et al., 2019). The use of nitrogen- and carbon-labeled amino acids provides more detailed information about proteins. Since the backbone of proteins is primarily composed of nitrogen and carbon atoms, ^15^N and ^13^C NMR data allow for the identification of which atoms belong to the main chain. Determining the interactions between backbone atoms is crucial for resolving secondary and tertiary structures (Wüthrich, 1990). As the size of the protein increases, the number of resonating atoms and, consequently, the amount of signal collected increases, making it more challenging to determine the positions of backbone atoms (Wüthrich, 1990). One of the reasons for this is that both alpha and beta carbons on the protein backbone produce NMR signals (Robson et al., 2018). The atom attached to the radical group is referred to as the alpha carbon, while the carbon belonging to the carboxyl group is called the beta carbon. The backbone of proteins consists of nitrogen from the amino group, the alpha carbon, and the beta carbon. When both carbons are NMR-active, the resonance of one atom affects the other, reducing the sensitivity of the measurement. This issue is resolved by producing proteins with ^13^C-labeled alpha carbons and ^12^C-labeled beta carbons (Robson et al., 2018).

In the standard production process using M9 minimal medium, glucose is utilized as the carbon source. Glucose serves both as an energy source and as a carbon donor for the synthesis of amino acids. However, the use of labeled glucose does not allow for selective labeling of amino acids with ^13^C. The use of labeled pyruvate as a carbon source, instead of labeled glucose, has been reported in the literature (Robson et al., 2018), enabling selective labeling of only the alpha carbons of amino acids. Some amino acids are synthesized directly from pyruvate, while others are produced through the gluconeogenesis pathway or the Krebs cycle. In all cases, the main carbon backbone of pyruvate is largely preserved, and the carbon atoms at specific positions are incorporated as alpha or beta carbons in the structure of amino acids. By using pyruvate labeled at the second or third carbon position, only the alpha carbons of amino acids can be selectively labeled, allowing for the collection of NMR data with higher sensitivity (Robson et al., 2018). Furthermore, this approach also simplifies resonance assignments.

While pyruvate-based isotopically labeled protein production is an effective method for resonance assignments, it also has some drawbacks. Compared to nutrient-rich media, the amount of protein produced is significantly lower, leading to a higher cost per unit of protein.

The minimal medium, which contains only various salts, a single nitrogen source, and a single carbon source, slows down bacterial metabolism, reducing the amount of protein produced. When pyruvate is used as the carbon source, certain amino acids are synthesized through the reversal of the glycolysis pathway. Since glycolysis operates in reverse to synthesize amino acids, a pathway typically used for energy production, this reversal negatively impacts bacterial metabolism and slows down protein production. Additionally, isotopically labeled chemicals are expensive due to the high production costs. As a result of these combined factors, the use of minimal medium and pyruvate as the carbon source results in a disproportionately high cost relative to the amount of protein produced. Therefore, there is a need to improve the protein production process when using pyruvate as the carbon source.

In bacterial cells, the regulation of intracellular metabolite levels is essential for maintaining metabolic balance and cellular homeostasis. Excess accumulation of specific metabolites can disrupt cellular processes and impede efficient metabolism. To avoid such disruptions, bacteria employ various mechanisms to expel unnecessary or excessive metabolites from the cell. For instance, waste products generated during metabolic processes, such as lactate, ethanol, or acetate, are often transported out of the cell to prevent intracellular buildup. Additionally, when certain metabolites, such as amino acids or organic acids, accumulate in large amounts, they can be exported to restore metabolic equilibrium. This export is mediated by specialized transporter proteins located in the bacterial membrane, which facilitate the active or passive transport of metabolites out of the cell. Efflux pumps, commonly associated with the removal of toxic compounds and antibiotics, also play a role in expelling surplus or unwanted metabolites. These transport and efflux systems are vital for bacterial survival, as they help preserve cellular homeostasis, optimize metabolic efficiency, and mitigate the toxic effects of accumulated byproducts. Understanding these mechanisms is critical for insights into bacterial metabolism and potential applications in biotechnology and medicine.

While pyruvate has been explored as an alternative carbon source for isotopic labeling in protein production, its impact on the overall metabolic transformations in bacterial culture medium remains under-investigated. Understanding how pyruvate influences metabolite composition and bacterial metabolism is critical for optimizing culture conditions and improving the efficiency of protein production. In this study, we aim to systematically investigate the metabolic changes in bacterial culture medium when pyruvate is used as the sole carbon source. By analyzing the metabolic shifts and metabolite profiles in the medium, we provide new insights into how pyruvate drives bacterial metabolic pathways, which can have broader implications for biotechnological processes and isotope labeling strategies.

## Material and Methods

### The transformation of the plasmid containing the target gene into the expression strain

In the first step, the transformation of GST fusion protein-tagged UbcH8 on the PGEX vector into *E. coli* BL21(DE3) cells has been carried out. For this, frozen bacterial cells and tubes containing plasmids have been placed on ice. The bacterial cells stored at -80°C and the plasmids stored at - 20°C have been allowed to reach the appropriate temperature. Lyophilized plasmids have first been centrifuged at 2650g for 3 minutes, and subsequently, 50 µL of ultra-pure water has been added. After vortexing for 30 seconds, they have been centrifuged again at 10600g for 4 minutes. The vortexing and centrifugation steps have been repeated, after which the plasmids have become ready for use. For the next step, a heat block has been set to 42°C, and sterilized 1.5mL Eppendorf tubes have been prepared. Into each tube, 50µL of thawed bacterial cells has been transferred. Similarly, 2µL of plasmid has been added to all tubes. The tubes have then been incubated on ice for 20 minutes. Following this, the tubes have been placed in a 42°C heat block for 45 seconds and immediately returned to ice. After 2 minutes, 500µL of pre-prepared, sterilized, room-temperature LB medium has been added to each tube. The tubes containing LB medium have been incubated at 37°C for 90 minutes with gentle inversion every 15 minutes. Subsequently, solid media plates containing ampicillin and chloramphenicol have been prepared for cell growth. The cells incubated for 90 minutes have been centrifuged at 238g for 2 minutes. The supernatant has been discarded, and the remaining cells in a small volume of liquid have been resuspended using a pipette. After the workspace has been sterilized with UV light, 50µL of cell suspension has been added to each agar plate, and the cells have been spread using glass beads. Once this step has been completed, the plates have been inverted and incubated at 37°C for 24 hours. For the next step, 150mL of sterilized LB medium in 250mL Erlenmeyer flasks has been supplemented with 150µL each of ampicillin and chloramphenicol solutions. Colonies from the solid cell cultures grown for 24 hours have been picked using pipette tips and transferred to the liquid culture. The liquid cultures have been placed in a shaking incubator set to 37°C and 110 rpm for 24 hours. Sterile, cryogenic tubes suitable for low-temperature storage have been prepared by adding 750µL of sterile glycerol. Into each tube, 750µL of the cell culture has been added, and the tubes have been sealed and stored in a -80°C freezer. The prepared transformed cells are now ready for use in initiating cell cultures when needed.

### Preparation of Pyruvate-based M9 Medium and Bacterial Growth

The next step after transformation for protein expression is the initiation of cell culture, which has been completed. For this, LB medium has been prepared in sterilized Erlenmeyer flasks. To ensure rapid bacterial growth, a rich medium has been used when starting the culture. LB powder, at a concentration of 25 grams per liter, has been dissolved in distilled water. Subsequently, 10 mL of the solution has been transferred into a flask. The flask has been sealed with aluminum foil and sterilized using an autoclave at 120 °C for 30 minutes under high pressure and steam. After cooling to room temperature, antibiotics (ampicillin and chloramphenicol) to which the transformed cells are resistant have been added to the flask. The transformed bacteria have been taken from the glycerol stock and transferred into the flask. The flask has been resealed and placed in an incubator set at 37 °C, allowing the bacteria to grow for at least 15 hours.

Before transferring the bacteria from the starter culture, a minimal medium containing unlabeled pyruvate has been prepared. As labeled and unlabeled pyruvate have identical chemical properties, normal pyruvate has been used to reduce experiment costs. For this, 0.3 grams of pyruvate have been weighed and dissolved in 100 mL of distilled water. Then, 0.43 grams of Na_2_HPO_4_, 0.36 grams of NaH_2_PO_4_, and 0.3 grams of KH_2_PO_4_ have been added (Robson et al., 2018). After adjusting the solution’s pH to 7 using NaOH, 0.1 grams of NH_4_Cl, 0.1 grams of NaHCO_3_, 0.024 grams of MgSO_4_, 0.0011 grams of CaCl_2_, and antibiotics (ampicillin and chloramphenicol) have been added. The medium has been sterilized using vacuum filtration (Robson et al., 2018).

From the bacterial culture grown in LB medium, 10 mL has been taken and centrifuged at 3000 rpm for 3 minutes to pellet the cells. The supernatant has been discarded, and 10 mL of the pre-prepared pyruvate-containing minimal medium has been added to the tube. The cells have been resuspended by pipetting, and the bacteria have been allowed to grow for 6 to 8 hours at 37°C, enabling adaptation from the rich medium to the minimal medium (Robson et al., 2018). The cells have then been centrifuged again at 3000 rpm for 3 minutes, and the supernatant has been replaced with fresh minimal medium. Samples have been taken from the culture hourly, and optical density has been measured.

When the optical density, measured at 595 and 600 nm wavelengths using a spectrophotometer, has reached 0.3, samples have been taken for NMR analysis. The incubator temperature has been lowered from 37 °C to 22 °C, as proteins tend to fold more correctly at lower temperatures (Rosano & Ceccarelli, 2014). When the optical density has reached 0.6, another sample has been collected. At an optical density of 0.6, IPTG has been added to induce protein production. Protein expression has been carried out for 15 hours, with optical density measurements taken during this time. All samples have been centrifuged immediately after collection to separate the medium from the cells. The samples have then been stored at -80 °C until NMR analysis could be performed.

### NMR sample preparation

After thawing the frozen medium, 500 µL was taken, and 55 µL of a buffer solution prepared with deuterium oxide was added. The added solution contained 37.7 mM Na2HPO4, 12.3 mM KH2PO4, 20 mM NaCl, and 1 mM DSS. DSS was included as a reference for the NMR analyses, and therefore it was added to achieve a final concentration of 0.1 mM. The prepared samples were placed in 5 mm NMR tubes, and measurements were performed. The NMR data acquisition was carried out using a 500 MHz Bruker Ascend magnet, equipped with an Avance NEO console and a BBO double resonance room temperature probe. In NMR-based metabolomics studies, water suppression techniques are typically employed to reduce the water peak rather than using a standard 1D proton pulse (Dag et al., 2022). For this study, we applied the 1D NOESY-presat (noesygppr1d) pulse sequence for all NMR data collections, with each spectrum consisting of 8K scans and 32K complex data points, over a spectral width of 9615.4 Hz. Chenomx NMR Suite 9.02 software (Chenomx Inc., Canada) was utilized to calculate the concentrations of target metabolites (Dag et al., 2022). Metabolite concentrations and the peak shapes of other compounds were determined by Chenomx, based on the concentration and peak height of DSS. The NMR data were collected with 8K scans to enhance the signal-to-noise ratio.

To analyze the NMR-based metabolomics data, we utilized MetaboAnalyst 6.0, an online platform designed for comprehensive metabolomics data processing and statistical analysis. Prior to statistical evaluations, the dataset was pre-processed using cube root transformation and Pareto scaling to normalize the data and reduce the influence of large variations among metabolites. For unsupervised exploration of the dataset, we applied Principal Component Analysis (PCA) to reduce the dimensionality of the data while capturing the maximum variance between samples. PCA allowed us to visualize the overall metabolic changes across different time points, revealing distinct clusters associated with the progression of bacterial growth and recombinant protein production, particularly in response to IPTG induction. In addition, Partial Least Squares Discriminant Analysis (PLS-DA) was employed as a supervised method to further discriminate between the time points and highlight key metabolites driving the separation. PLS-DA is a powerful statistical tool that maximizes the separation between predefined groups, in this case, different time points during the growth and production phases. The Variable Importance in Projection (VIP) scores obtained from PLS-DA identified the most influential metabolites contributing to the temporal metabolic changes observed in the bacterial culture. Finally, Pearson correlation analysis was performed to assess the relationships between individual metabolites, followed by hierarchical clustering to group metabolites based on their correlation patterns. These analyses revealed distinct clusters of metabolites that are co-regulated, providing insights into the metabolic pathways that are modulated during recombinant protein production.

## Results and Discussion

### Bacterial Growth in Pyruvate-Based Media

The optical density (OD) measurements at 600 nm were taken at various time points to assess the bacterial growth in three separate flasks (Figure 1). At the start of the experiment (0 hours), the initial OD values were low, with Flask 1 at 0.108, Flask 2 at 0.090, and Flask 3 at 0.091, indicating a low bacterial density. After 3 hours, the OD values increased slightly, with Flask 1 measuring 0.23, Flask 2 at 0.223, and Flask 3 at 0.229, demonstrating early bacterial growth. At 9 hours, just prior to the addition of IPTG for protein induction, the OD values had further increased to approximately 0.55 in all flasks. Following IPTG induction at 9 hours, the OD values at 10 hours (1 hour post-induction) showed a small but consistent rise, with values around 0.55 in all flasks. By 24 hours (14 hours post-induction), the cultures reached their peak OD readings, with Flask 1 at 0.742, Flask 2 at 0.782, and Flask 3 at 0.772. At 26 hours (16 hours post-induction), the OD values remained stable around 0.78 for all flasks, indicating continued growth. However, at the 36-hour time point (26 hours post-induction), a slight decrease in OD was observed across all flasks, with values dropping to 0.679 in Flask 1, 0.667 in Flask 2, and 0.700 in Flask 3. This decline suggests that the bacterial cultures may have entered the stationary phase or experienced nutrient depletion in the medium.

**Figure 1.**
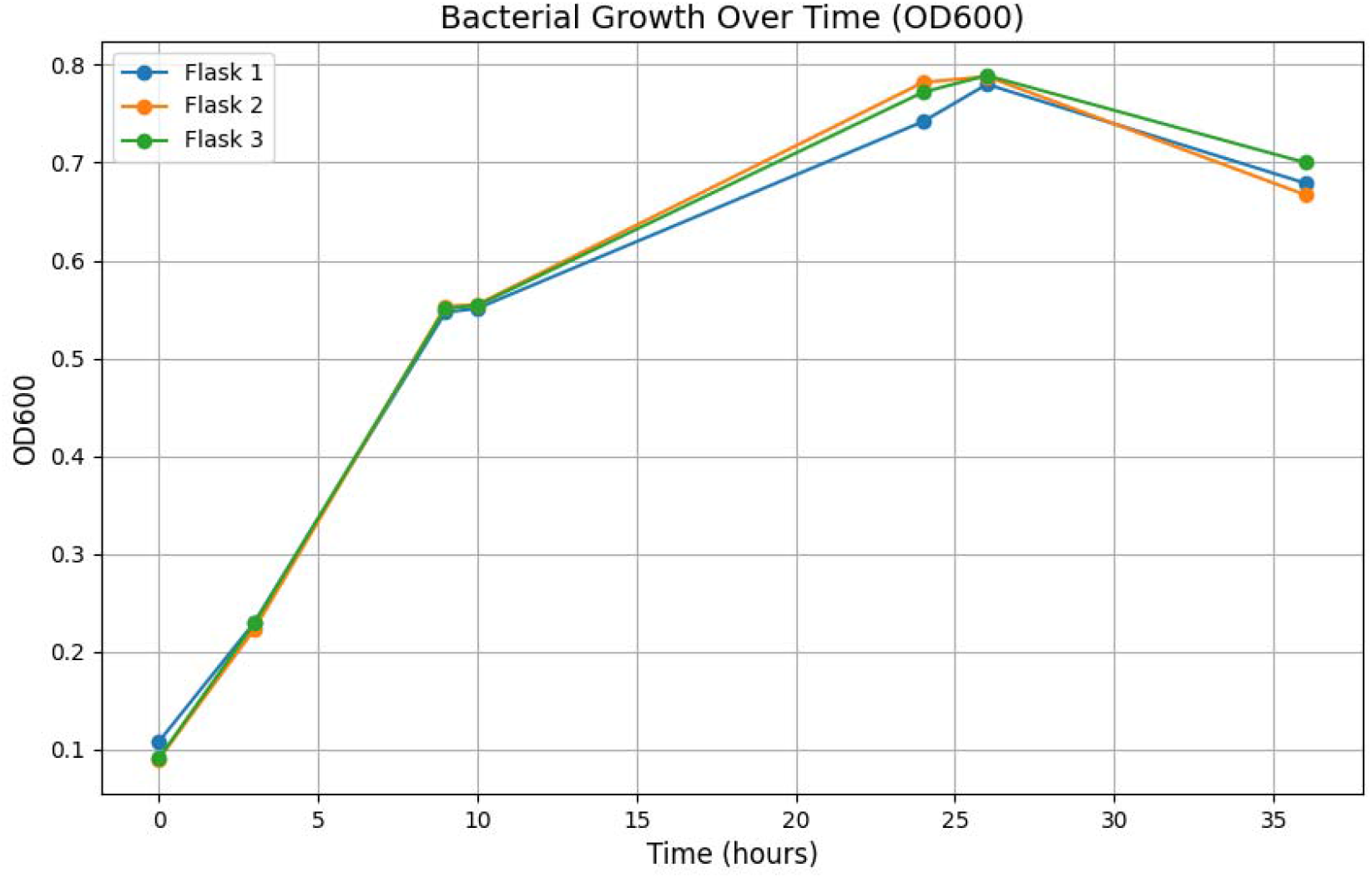
Comparative Analysis of Bacterial Growth in Three Independent Cultures Over 36 Hours, Monitored by Optical Density at 600 nm

### Metabolite Profiling in the Culture Medium

The bar plots in Figure 2 illustrate the temporal concentration changes of key metabolites involved in central metabolic pathways during bacterial growth and recombinant protein production. These metabolites were measured at various time points: 0h, 3h, 9h (IPTG induction), 10h, 24h, 26h, and 36h, capturing metabolic dynamics from the lag phase through the exponential and stationary phases of bacterial growth. The analysis of metabolite concentrations across different time points revealed significant changes in key intermediates involved in central carbon metabolism during bacterial growth and recombinant protein production. These metabolic shifts were observed both before and after IPTG induction, reflecting the cells’ metabolic response to the high availability of pyruvate as the primary carbon source.

**Figure 2.**
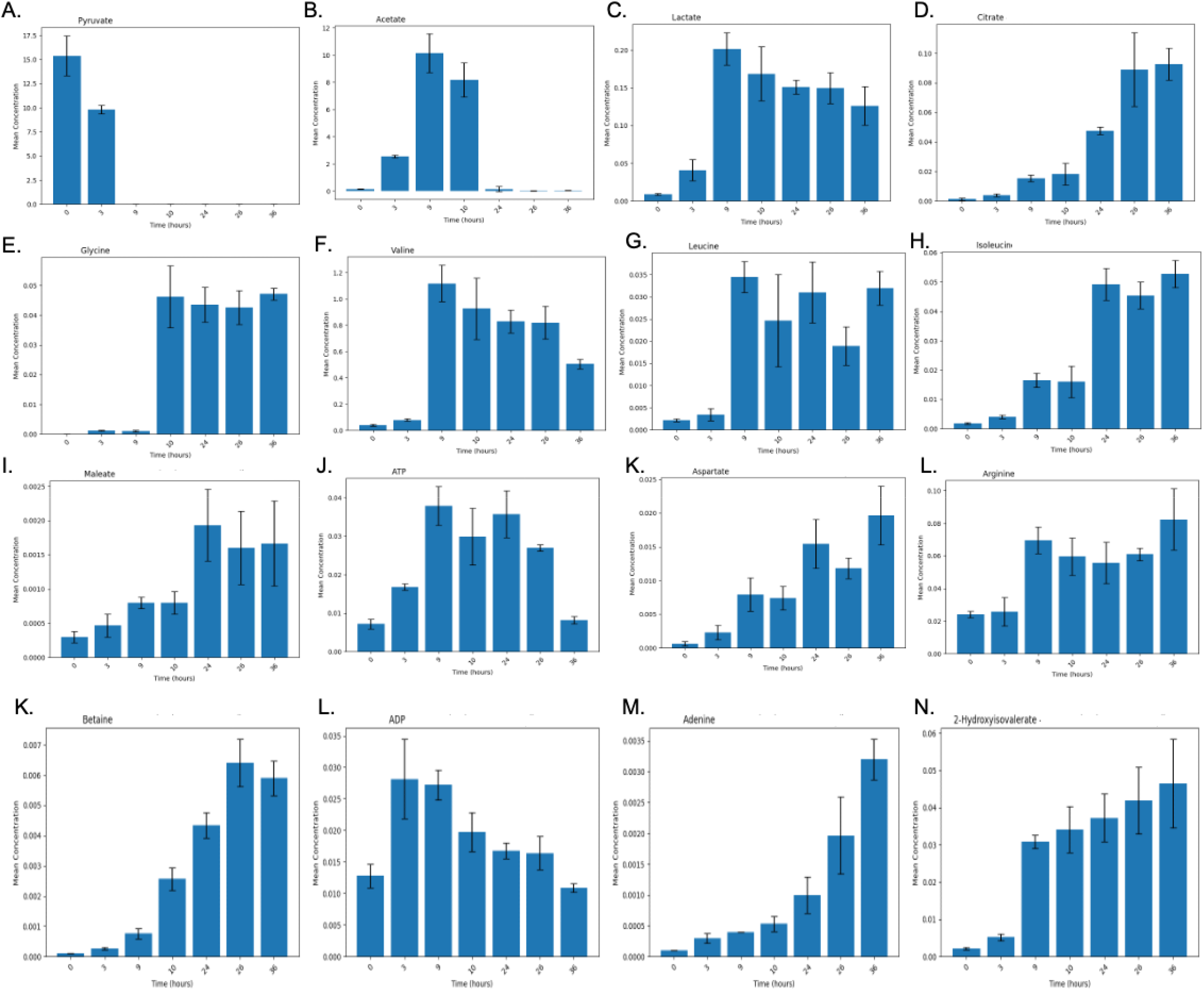
Time-Dependent Concentration Profiles of Key Metabolites during Bacterial Growth and Recombinant Protein Production. A. Pyruvate, B. Acetate, C. Lactate, D. Citrate, E. Glycine, F. Valine, G. Leucine, H. Isoleucine, I. Maleate, J. ATP, K. Aspartate, L. Arginine, M. Adenine, N. 2-Hydroxyisovalerate.

Pyruvate (Figure 2A), the main carbon source in this experiment, exhibited a rapid decrease starting well before IPTG induction, with its levels already substantially reduced by the 9-hour mark. This early decline in pyruvate suggests that bacterial cells are actively consuming pyruvate through multiple metabolic pathways to support energy production and biosynthesis. The sustained reduction of pyruvate post-9h further indicates that the cells continue to use pyruvate intensively during recombinant protein production. The early depletion of pyruvate is likely a result of its diversion into multiple pathways, including both the TCA cycle for energy production and overflow metabolic pathways. Concurrent with the depletion of pyruvate, both acetate (Figure 2B) and lactate (Figure 2C) showed substantial increases in the culture media prior to IPTG induction. This observation suggests that bacterial cells, faced with an abundance of pyruvate, are converting excess pyruvate into acetate and lactate as a means of mitigating the buildup of intracellular pyruvate. These metabolites are likely excreted into the medium as waste products to avoid toxic intracellular accumulation. The high concentrations of acetate and lactate in the culture media reflect overflow metabolism, a process whereby excess carbon from glycolysis (in the form of pyruvate) is diverted into pathways that produce acetate and lactate, which are then secreted from the cells. This process is typical of bacterial cultures experiencing rapid growth and high glycolytic flux, where the TCA cycle becomes saturated and alternative metabolic routes are used to handle the surplus carbon.

After IPTG induction at 9 hours, the concentrations of acetate and lactate in the medium remained elevated. The sustained high levels of acetate and lactate post-IPTG likely reflect the fact that once these metabolites are expelled into the culture medium, their reuptake and reintegration into intracellular metabolism is limited. Acetate and lactate, once secreted, may no longer serve as useful substrates for energy production or biosynthesis. Instead, their accumulation in the medium suggests that the bacterial cells are unable, or inefficient, at reclaiming these byproducts. Citrate (Figure 2D), a key intermediate of the TCA cycle, showed a gradual increase in concentration, particularly after IPTG induction. This suggests that while pyruvate is being converted into acetate and lactate, a portion of it continues to fuel the TCA cycle to meet the energy demands and biosynthetic needs of the cells. Citrate’s rise is indicative of active TCA cycle flux, particularly after recombinant protein production begins, as the cells require sustained energy and carbon skeletons for the synthesis of amino acids and other macromolecules. Together, the patterns observed in pyruvate, acetate, lactate, and citrate concentrations highlight the metabolic flexibility of bacterial cells in response to pyruvate availability. Before IPTG induction, excess pyruvate is channeled into overflow pathways, resulting in the secretion of acetate and lactate into the medium. Following IPTG induction, this pattern continues, but the rise in citrate suggests that a portion of the pyruvate is also directed into the TCA cycle to support the increased metabolic demands of recombinant protein production. The secretion of acetate and lactate into the culture media throughout the experiment underscores the cells’ need to expel excess metabolites and maintain intracellular metabolic balance under conditions of high pyruvate flux.

Glycine (Figure 2E) shows stable concentrations before IPTG induction, followed by a modest increase during the later time points. This suggests that glycine metabolism is linked to protein biosynthesis during the extended growth phases. Branched-chain amino acids (BCAAs), including valine (Figure 2F), leucine (Figure 2G), and isoleucine (Figure 2H), show moderate increases post-IPTG induction, indicative of their essential roles in protein synthesis. The upregulation of BCAA biosynthesis following IPTG induction reflects the heightened demand for amino acids during recombinant protein production. The concentration of maleate (Figure 2I) increases gradually, peaking at 24 hours, suggesting its involvement in energy production pathways as bacterial cells transition into the stationary phase. ATP (Figure 2J) levels rise significantly after IPTG induction, peaking at 10 hours, indicating an increased demand for energy during the early stages of recombinant protein synthesis. This is followed by a gradual decline in ATP levels, reflecting a stabilization of energy metabolism as the bacterial culture approaches stationary phase. The ADP (Figure 2L) profile mirrors that of ATP, peaking at 10 hours and gradually decreasing, indicating high energy turnover during protein production.

Aspartate (Figure 2K) and arginine (Figure 2L) show continuous increases in concentration after IPTG induction, with aspartate peaking at 24 hours. This suggests their critical roles in supporting protein synthesis and nitrogen metabolism, particularly as amino acid demand rises during recombinant protein production. Similarly, betaine (Figure 2K) concentrations increase over time, peaking at 24 hours. Betaine is known to function as an osmoprotectant, and its accumulation may reflect bacterial stress responses during extended periods of growth and protein synthesis. Adenine (Figure 2M), a purine nucleotide, increases steadily following IPTG induction, peaking at 36 hours, indicating heightened nucleotide turnover associated with increased DNA/RNA synthesis during recombinant protein production. Finally, 2- hydroxyisovalerate (Figure 2N), a metabolite linked to branched-chain amino acid metabolism, shows a gradual increase over time, peaking at 36 hours. This suggests its involvement in secondary metabolic pathways, particularly as bacterial cells shift toward the stationary phase and metabolic processes become focused on maintenance rather than growth.

Taken together, these results reveal a clear metabolic reprogramming in response to IPTG induction and recombinant protein production. Pyruvate consumption, acetate overflow, and shifts in amino acid and nucleotide metabolism reflect the bacterial cells’ need to balance energy production with the synthesis of macromolecules for protein production. The most significant metabolic changes occur between 9 and 10 hours, shortly after IPTG induction, indicating the critical metabolic adjustments required to meet the demands of protein synthesis. As the bacterial culture transitions to the stationary phase (24-36 hours), metabolic activity stabilizes, as evidenced by the gradual leveling off of key metabolite concentrations.

### Multivariate Analysis of Metabolite Profiles

The PCA scores plot (Figure 3A) highlights distinct separation of metabolite profiles across time points, with the first two principal components (PC1 and PC2) explaining 49.9% and 22.4% of the total variance, respectively. The samples collected at 0h and 3h cluster tightly, indicating minimal metabolic variation during the early growth (lag) phase when bacterial metabolism is primarily focused on adjusting to the medium. However, a significant shift occurs between 3h and 9h, corresponding to the addition of IPTG at the 9-hour mark to induce recombinant protein production.

**Figure 3.**
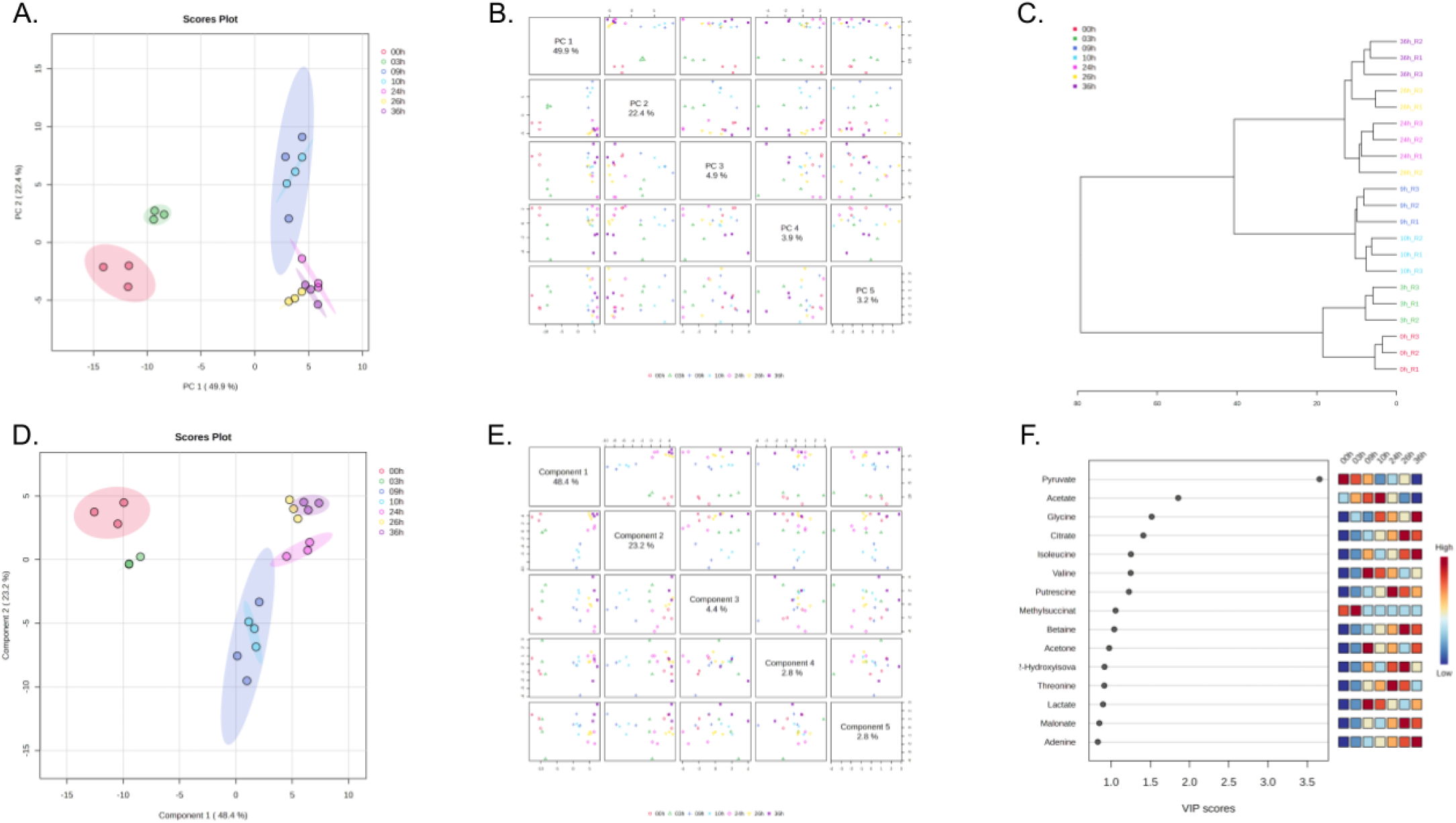
Multivariate Analysis of Metabolite Profiles at Different Time Points. **A. Principal Component Analysis (PCA) Scores Plot** showing the separation of metabolite profiles at various time points (0h, 3h, 9h, 10h, 24h, 26h, and 36h). The PCA demonstrates distinct clustering of samples from different time points, with PC1 explaining 49.9% of the variance and PC2 explaining 22.4%. **B. Pairwise Plot of Principal Components (PCs)**, illustrating the relationships between different PCs from the PCA analysis. **C. Hierarchical Clustering Dendrogram** visualizing the similarity of metabolite profiles across replicates at different time points, indicating clear time-dependent grouping of samples. **D. Partial Least Squares Discriminant Analysis (PLS-DA) Scores Plot** highlighting the discriminative power of PLS-DA, with Component 1 explaining 48.4% of the variance and Component 2 explaining 22.3%. The plot shows distinct separation of metabolic profiles over time, similar to the PCA results. **E. Pairwise Plot of PLS-DA Components** showing the relationship between components generated by the PLS-DA model. **F. Variable Importance in Projection (VIP) Scores** from the PLS-DA model, ranking metabolites based on their contribution to class separation. Metabolites such as acetate, glycine, and citrate are the top contributors to the separation between time points, with corresponding heat maps showing their concentration changes over time.

Post-induction, the 10h samples (1 hour after IPTG induction) show a distinct shift along PC1, indicating rapid metabolic changes following protein induction. This is likely due to the metabolic burden of recombinant protein synthesis, which alters the demand for energy and metabolic intermediates derived from pyruvate. Notably, the 24h and 26h time points (14 and 16 hours after IPTG induction, respectively) form closely clustered groups, suggesting that the bacteria have reached a more stable metabolic state during this extended period of protein production. This clustering is consistent with the bacteria entering the stationary phase, during which growth slows and metabolism stabilizes, with a potential shift toward metabolite recycling and waste product excretion (e.g., acetate, lactate).

By 36h (26 hours post-induction), the metabolite profile has shifted slightly compared to earlier post-IPTG time points, likely due to nutrient depletion or accumulation of metabolic byproducts in the culture medium. The confidence ellipses show tight clustering of replicate samples at later time points (24h, 26h, 36h), indicating reproducibility of metabolic states during and after recombinant protein production. The PCA results clearly indicate that IPTG induction at 9h causes a dramatic metabolic shift, with the most pronounced changes occurring between 9h and 10h. These shifts reflect the metabolic adaptations required for the bacteria to support increased protein synthesis, including changes in energy metabolism, amino acid synthesis, and metabolite excretion. The pairwise scatter plots of principal components (Figure 3B) provide a more detailed view of the variance captured by the top five components. The first two principal components (PC1 and PC2) capture most of the variance, as indicated by the clear clustering of samples according to time points, especially around IPTG induction (9h-10h). Time points immediately following IPTG induction, such as 10h, show clear separation from pre-induction time points, illustrating the significant metabolic impact of initiating recombinant protein synthesis. Minor components, such as PC3 (4.9%) and PC4 (3.9%), capture additional subtle metabolic changes that may reflect variations in the levels of specific metabolites involved in later phases of growth and protein production. For instance, these components might be influenced by the accumulation of waste products like acetate or lactate as bacterial growth slows. In summary, the PCA analysis reveals significant metabolic shifts corresponding to the addition of IPTG at 9h, with marked differences in metabolite profiles between pre- and post-induction phases. The clustering of time points from 24h to 36h suggests that bacterial metabolism stabilizes during the later stages of protein production, possibly due to nutrient limitation or the plateauing of recombinant protein synthesis.

To further investigate the relationships between the metabolite profiles at different time points during recombinant protein production, a hierarchical clustering analysis was performed (Figure 3C). The clustering dendrogram groups samples based on their metabolic similarity, revealing clear temporal patterns in the data. The samples collected at 0h, 3h, and 9h (prior to or at the time of IPTG induction) form a distinct cluster, indicating that the metabolic profiles during the early phases of growth (lag and early exponential phases) are highly similar. The proximity of these time points suggests minimal metabolic reprogramming in the absence of IPTG and during the initial phase of bacterial growth in the pyruvate-supplemented medium. The 9h time point represents the moment IPTG is added to induce recombinant protein production, yet it remains metabolically similar to earlier time points, reflecting a delay in the metabolic response to IPTG. In contrast, the samples collected 1 hour after IPTG induction (10h) form a separate, highly distinct cluster from the earlier time points. This separation underscores the significant metabolic changes that occur rapidly following IPTG induction, likely driven by the metabolic demands associated with recombinant protein synthesis. These changes may include shifts in energy metabolism, amino acid biosynthesis, and central carbon metabolism, all necessary to support the increased biosynthetic activity. The 24h, 26h, and 36h time points (14, 16, and 26 hours post-IPTG induction, respectively) form a tight, well-defined cluster. This clustering indicates that once the bacteria have adapted to the protein production process, their metabolism stabilizes. The minimal differences between these time points suggest that bacterial cells have reached a stationary phase, with metabolic activities largely focused on maintaining cellular homeostasis and recycling metabolites. This stabilization is likely due to nutrient depletion and the plateauing of protein synthesis as bacterial growth slows down. The close grouping of replicates within these time points further demonstrates the reproducibility of the metabolic changes observed.

Interestingly, hierarchical clustering also reveals that some of the 9h replicates show partial overlap with 10h replicates, suggesting a transitional phase in metabolic reprogramming as cells switch from regular growth to recombinant protein production. This gradual transition is expected, as the induction of recombinant protein synthesis imposes a substantial metabolic burden, requiring time for bacterial cells to adjust their metabolic pathways to meet the demands of protein synthesis. Overall, the hierarchical clustering analysis provides a clear temporal overview of the metabolic shifts that occur during bacterial growth and recombinant protein production. The clustering of time points prior to IPTG induction, followed by distinct metabolic states post-induction, highlights the significant metabolic reprogramming that accompanies protein production. This analysis confirms that bacterial metabolism stabilizes during the later stages of growth, likely in response to nutrient limitation and the completion of protein synthesis. To complement the unsupervised analysis provided by PCA, Partial Least Squares Discriminant Analysis (PLS-DA) was performed to further explore the metabolic changes in response to IPTG induction and identify metabolites that contribute most to the observed variations across time points. The PLS-DA scores plot (Figure 3D) highlights the discriminative power of the PLS-DA model, which maximizes the separation between metabolite profiles at different time points. In this plot, Component 1 explains 48.4% of the variance, and Component 2 explains 22.3%, together capturing a significant portion of the total variability. The clear separation of the time points along these two components reflects the dynamic changes in bacterial metabolism over time, particularly in response to IPTG induction at the 9-hour mark. Pre-induction time points, 0h, 3h, and 9h, form distinct clusters, with 9h showing more separation from earlier time points, as expected, given that IPTG induction occurs at this time. Interestingly, the 9h and 10h samples (1 hour post-IPTG induction) show a sharp metabolic shift, as these two clusters are well separated, indicating a rapid metabolic reprogramming following IPTG addition. This shift is likely due to the metabolic demands imposed by recombinant protein production, which alters the cellular utilization of pyruvate and other metabolites involved in central carbon metabolism and amino acid biosynthesis. The later time points, 24h, 26h, and 36h, form tightly clustered groups, with some overlap, indicating that the bacterial metabolism reaches a more stable state after several hours of protein production. This stabilization likely reflects the adaptation of bacterial cells to the metabolic burden of protein production and the onset of the stationary phase, where growth has slowed, and nutrient consumption is reduced. The tighter clustering at these time points suggests that while metabolic activity persists, the variation between these samples is minimal, consistent with a plateau in protein synthesis and bacterial growth. The confidence ellipses around each time point cluster indicate the consistency of replicate samples, particularly for the post-induction time points, showing that the metabolic state of the cultures is highly reproducible after the initial response to IPTG. This distinct separation of time points in the PLS-DA plot, particularly around the 9h and 10h time points, confirms that IPTG induction is a major driver of metabolic variation in the dataset. The pairwise scatter plots of the first five components of the PLS-DA model (Figure 3E) provide a more detailed breakdown of how each component contributes to the separation of time points. As shown in the plot, Component 1(48.4%) and Component 2 (22.3%) account for most of the variance, with clear separation of early (0h, 3h) and post-induction time points. This further supports the observation that IPTG induction introduces significant metabolic shifts, which are well captured by the first two components. Other components, such as Component 3 (4.4%) and Component 4 (2.8%), capture more subtle variations that may be related to secondary metabolic processes or changes in less abundant metabolites that contribute to the overall metabolic profile. These components likely reflect more nuanced variations in the cellular metabolism, such as changes in specific amino acid levels or byproduct excretion (e.g., acetate, lactate) as the bacteria adapt to prolonged recombinant protein production. The pairwise component plots confirm that the metabolic changes induced by IPTG are most strongly captured by Components 1 and 2, with minimal variance explained by subsequent components. This suggests that the major metabolic reprogramming in response to IPTG induction can be largely described by the shifts along these two axes, with smaller components reflecting minor metabolic adjustments as the cultures reach the stationary phase.The PLS-DA analysis provides a supervised view of the temporal metabolic shifts in bacterial cultures during recombinant protein production. The analysis confirms that IPTG induction at 9h is a major driver of metabolic variation, leading to rapid and distinct metabolic changes as bacteria adapt to the demands of protein synthesis. The stabilization of metabolite profiles at later time points, as seen in both the scores plot and pairwise component plots, reflects the transition into the stationary phase, where nutrient consumption slows, and the metabolic state becomes more consistent.

To further understand the metabolites driving the separation of time points in the PLS-DA analysis, we calculated the Variable Importance in Projection (VIP) scores (Figure 3F). VIP scores rank metabolites based on their contribution to the PLS-DA model, identifying those most influential in distinguishing between time points, particularly in response to IPTG induction and the subsequent phases of bacterial growth and protein production. The metabolites with the highest VIP scores are displayed in Figure 3F, indicating their importance in explaining the metabolic shifts across the time points. The top-ranking metabolites include pyruvate, acetate, glycine, citrate, isoleucine, and valine, which are directly or indirectly involved in energy metabolism, amino acid biosynthesis, and central carbon metabolism. These metabolites are particularly important in the context of bacterial growth under minimal medium conditions, where pyruvate serves as the sole carbon source.

Pyruvate, the primary carbon source used in this study, ranks as the most important metabolite, as expected. Pyruvate is not only a key energy source but also a central intermediate in multiple metabolic pathways, including the citric acid cycle (TCA cycle) and gluconeogenesis. Its importance reflects the fact that the metabolic shifts occurring after IPTG induction and during recombinant protein production are tightly linked to how pyruvate is utilized and converted into essential biomolecules. Acetate, a common byproduct of bacterial metabolism, also ranks highly in the VIP score. Its accumulation is typically a sign of overflow metabolism, where excess carbon from pyruvate is converted into acetate, especially under conditions where the TCA cycle is saturated or energy demands exceed the capacity of oxidative metabolism. This aligns with the later time points, where bacterial growth slows, and acetate may be secreted as a waste product during the stationary phase. Glycine, an amino acid involved in protein biosynthesis and the folate cycle, also contributes significantly to the separation of time points. Its importance in the VIP score suggests that shifts in glycine metabolism are closely linked to the demands of protein production, especially after IPTG induction, when the need for amino acids increases. Citrate, a key metabolite in the TCA cycle, plays a critical role in energy production and biosynthesis. The high VIP score for citrate indicates that variations in its levels are essential in distinguishing between time points, reflecting the dynamic shifts in energy metabolism as bacteria transition through different growth phases, particularly in response to the metabolic burden of recombinant protein synthesis. Isoleucine and valine, both branched-chain amino acids (BCAAs), are also among the top-ranked metabolites. These amino acids are not only essential for protein synthesis but are also linked to energy metabolism. Bacteria may upregulate BCAA biosynthesis or alter their catabolism in response to the increased demand for protein production following IPTG induction. The significant contribution of these amino acids to the PLS-DA model suggests that their levels are highly responsive to the metabolic state of the culture and the progression through growth phases. To complement the VIP score rankings, the heatmap on the right side of Figure 3F shows the relative concentrations of these key metabolites across the different time points. This visualization highlights the dynamic changes in metabolite levels over time, with warmer colors (red) indicating higher concentrations and cooler colors (blue) indicating lower concentrations. For example, pyruvate levels decrease after IPTG induction, suggesting its rapid utilization in metabolic pathways during protein production. In contrast, acetate levels increase over time, consistent with its role as a byproduct of overflow metabolism, especially as cells transition into the stationary phase. Glycine, citrate, and BCAAs (isoleucine, valine) also exhibit time-dependent fluctuations, reflecting their roles in protein synthesis and energy metabolism. The heatmap provides a clear visualization of how the concentration of these metabolites changes in response to bacterial growth and protein production. The most substantial shifts are seen post-IPTG induction, particularly at the 10h, 24h, and 36h time points, corresponding to the exponential and stationary phases of bacterial growth. The VIP score analysis identifies key metabolites that are critical in explaining the metabolic shifts observed during the bacterial growth phases and in response to IPTG induction. Pyruvate, acetate, glycine, citrate, and branched-chain amino acids play pivotal roles in supporting energy metabolism and protein biosynthesis under minimal medium conditions with pyruvate as the carbon source. The heatmap further illustrates the dynamic changes in these metabolites, providing insights into how bacterial metabolism adapts to the metabolic demands of recombinant protein production.

To understand the relationships between metabolites during recombinant protein production, we performed a Pearson correlation analysis, followed by hierarchical clustering to group metabolites based on their co-regulation patterns. These analyses provide insights into how metabolites interact with one another during different phases of bacterial growth. The heatmap in Figure 4A shows the Pearson correlation coefficients between metabolites across all time points, providing a comprehensive view of metabolite interactions. The color scale ranges from dark red (strong positive correlation, close to +1) to dark blue (strong negative correlation, close to -1), with white indicating no correlation. Several distinct clusters of highly correlated metabolites emerge, suggesting that these groups are co-regulated during bacterial growth and recombinant protein production. Pyruvate, a central metabolite in glycolysis and the citric acid cycle, shows strong positive correlations with several key energy-related metabolites such as acetate, succinate, and citrate, reflecting its pivotal role in energy metabolism and carbon flow through these pathways. The tight clustering of these metabolites suggests that they are part of a coordinated metabolic response to the use of pyruvate as the sole carbon source. In contrast, metabolites such as NADPH and NADH, which are involved in redox reactions and cellular energy balance, display negative correlations with certain amino acids like isoleucine and valine, possibly reflecting a metabolic trade-off between energy production and biosynthesis during recombinant protein production. This is consistent with the metabolic burden that bacteria face when diverting resources toward protein synthesis. The metabolites involved in amino acid metabolism, such as leucine, valine, and isoleucine (branched-chain amino acids, BCAAs), form a highly correlated cluster. These BCAAs play critical roles in protein synthesis and are tightly regulated during recombinant protein production, particularly after IPTG induction. Their strong correlations with other amino acids, such as serine and glycine, suggest that the demand for amino acids during the growth and production phases drives coordinated metabolic shifts.

**Figure 4.**
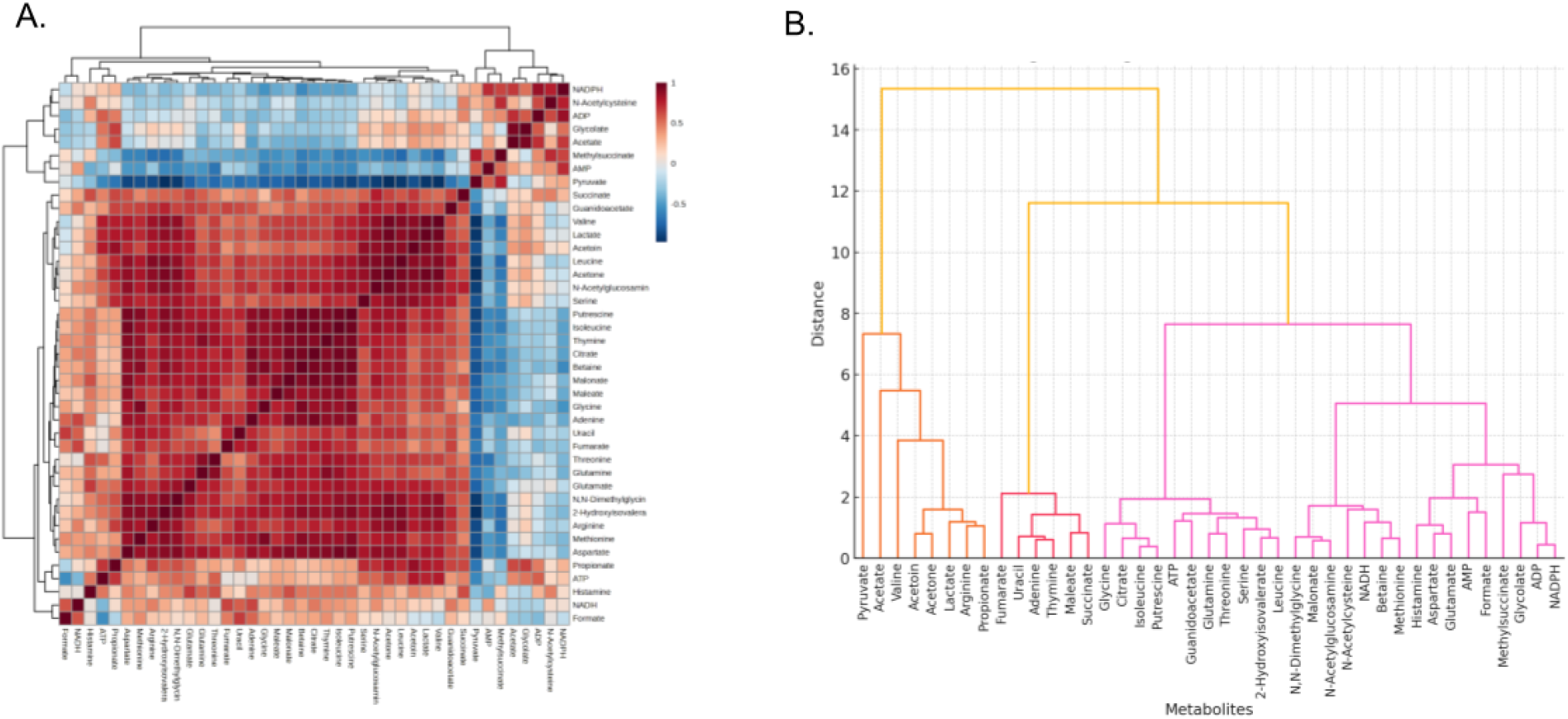
Correlation Matrix and Hierarchical Clustering of Metabolites in Bacterial Cultures during Recombinant Protein Production **A. Heatmap of Pearson Correlation Coefficients** between metabolites, illustrating the pairwise correlations of metabolite levels during different phases of bacterial growth and protein production. The color scale ranges from strong positive correlations (red) to strong negative correlations (blue). Metabolites involved in energy metabolism, amino acid biosynthesis, and central carbon metabolism form distinct clusters, reflecting their co-regulation during bacterial growth. **B. Hierarchical Clustering Dendrogram** showing the relationships between metabolites based on similarity in their correlation patterns. Metabolites are grouped into clusters based on their correlation distances, with closer distances indicating higher similarity in their metabolic roles. Key metabolites such as pyruvate, acetate, and branched-chain amino acids (BCAAs) are highlighted as major contributors to metabolic shifts during the recombinant protein production process.

**Table 1.** Concentration of Metabolites at Different Time Points during Bacterial Growth and Recombinant Protein Production. The concentrations of selected metabolites measured in the culture media at multiple time points (0h, 3h, 9h, 10h, 24h, 26h, and 36h). Metabolite concentrations are reported for three biological replicates (R1, R2, and R3) for each time point.

| Compound Name | 0h R1 | 0h R2 | 0h R3 | 3h R1 | 3h R2 | 3h R3 | 9h R1 | 9h R2 | 9h R3 | 10h R1 | 10h R2 | 10h R3 | 24h R1 | 24h R2 | 24h R3 | 26h R1 | 26h R2 | 26h R3 | 36h R1 | 36h R2 | 36h R3 |
| --- | --- | --- | --- | --- | --- | --- | --- | --- | --- | --- | --- | --- | --- | --- | --- | --- | --- | --- | --- | --- | --- |
| 2-Hydroxyisovalerate | 0.0017 | 0.0022 | 0.0026 | 0.0043 | 0.0049 | 0.0064 | 0.0311 | 0.0286 | 0.0329 | 0.027 | 0.0334 | 0.042 | 0.0294 | 0.0372 | 0.0452 | 0.0545 | 0.0348 | 0.0365 | 0.0355 | 0.063 | 0.0409 |
| Acetate | 0.1649 | 0.1207 | 0.13 | 2.4283 | 2.5641 | 2.6352 | 9.771 | 12.0149 | 8.5956 | 6.5189 | 8.4271 | 9.5573 | 0.4077 | 0.0121 | 0.0126 | 0.0151 | 0.0248 | 0.0206 | 0.0402 | 0.0579 | 0.04 |
| Acetoin | 0.009 | 0.0129 | 0.0101 | 0.0264 | 0.0243 | 0.0288 | 0.0516 | 0.0656 | 0.069 | 0.0336 | 0.0458 | 0.0589 | 0.041 | 0.0573 | 0.0548 | 0.0418 | 0.0395 | 0.0282 | 0.0332 | 0.0382 | 0.0452 |
| Acetone | 0.0018 | 0.0027 | 0.0052 | 0.0116 | 0.0153 | 0.0166 | 0.0939 | 0.1192 | 0.1012 | 0.0669 | 0.0878 | 0.099 | 0.0643 | 0.0795 | 0.0762 | 0.0684 | 0.078 | 0.0747 | 0.0818 | 0.0771 | 0.0848 |
| Adenine | 0.0001 | 0.0001 | 0.0001 | 0.0002 | 0.0003 | 0.0004 | 0.0004 | 0.0004 | 0.0004 | 0.0004 | 0.0005 | 0.0007 | 0.0006 | 0.0013 | 0.0011 | 0.0018 | 0.0028 | 0.0013 | 0.0028 | 0.0036 | 0.0032 |
| ADP | 0.0154 | 0.0109 | 0.0121 | 0.0193 | 0.0321 | 0.0331 | 0.024 | 0.0296 | 0.0281 | 0.0154 | 0.0215 | 0.0223 | 0.0184 | 0.0166 | 0.0153 | 0.0128 | 0.0189 | 0.0176 | 0.0116 | 0.0112 | 0.01 |
| AMP | 0.0185 | 0.0128 | 0.0107 | 0.0153 | 0.0174 | 0.014 | 0.0117 | 0.0143 | 0.0161 | 0.0089 | 0.0123 | 0.0105 | 0.0084 | 0.0086 | 0.0074 | 0.0067 | 0.0111 | 0.0091 | 0.0129 | 0.0114 | 0.0135 |
| Arginine | 0.0264 | 0.0242 | 0.0215 | 0.0208 | 0.0186 | 0.0381 | 0.0591 | 0.0792 | 0.0707 | 0.0491 | 0.0542 | 0.0755 | 0.0416 | 0.0722 | 0.0535 | 0.0557 | 0.0647 | 0.0626 | 0.0631 | 0.0763 | 0.1078 |
| Aspartate | 0.0002 | 0.0007 | 0.001 | 0.0009 | 0.0026 | 0.0034 | 0.0055 | 0.0113 | 0.007 | 0.0056 | 0.0068 | 0.0098 | 0.0122 | 0.0205 | 0.0136 | 0.0117 | 0.01 | 0.0138 | 0.02 | 0.0142 | 0.0248 |
| ATP | 0.0088 | 0.0056 | 0.0071 | 0.0178 | 0.0158 | 0.0169 | 0.0314 | 0.0437 | 0.0384 | 0.0202 | 0.0313 | 0.0381 | 0.0272 | 0.0408 | 0.0392 | 0.0266 | 0.0282 | 0.0263 | 0.0092 | 0.0069 | 0.0085 |
| Betaine | 0.0001 | 0.0001 | 0.0001 | 0.0002 | 0.0003 | 0.0003 | 0.0007 | 0.0006 | 0.001 | 0.0026 | 0.003 | 0.0021 | 0.0039 | 0.0049 | 0.0042 | 0.0073 | 0.0065 | 0.0054 | 0.0056 | 0.0067 | 0.0054 |
| Citrate | 0.0006 | 0.0009 | 0.0022 | 0.0028 | 0.0042 | 0.0049 | 0.0128 | 0.0155 | 0.018 | 0.0113 | 0.015 | 0.0284 | 0.0457 | 0.0511 | 0.0457 | 0.0552 | 0.1144 | 0.0976 | 0.0863 | 0.1079 | 0.084 |
| Formate | 0.0021 | 0.0023 | 0.0028 | 0.0029 | 0.0053 | 0.0122 | 0.0048 | 0.0002 | 0.0004 | 0.0034 | 0.0044 | 0.005 | 0.0025 | 0.0026 | 0.0036 | 0.0024 | 0.0019 | 0.0026 | 0.0162 | 0.0252 | 0.0202 |
| Glutamate | 0.0028 | 0.0045 | 0.0058 | 0.0121 | 0.0066 | 0.0103 | 0.0089 | 0.0169 | 0.0103 | 0.0098 | 0.0125 | 0.0137 | 0.0128 | 0.0148 | 0.0204 | 0.0173 | 0.021 | 0.0103 | 0.0136 | 0.0169 | 0.0159 |
| Glutamine | 0.0037 | 0.0063 | 0.0085 | 0.0125 | 0.0139 | 0.0176 | 0.0097 | 0.0257 | 0.0176 | 0.0144 | 0.0208 | 0.013 | 0.0398 | 0.0785 | 0.0644 | 0.0216 | 0.0337 | 0.0304 | 0.0309 | 0.065 | 0.0309 |
| Glycine | 0.0001 | 0.0001 | 0.0001 | 0.0009 | 0.0011 | 0.0014 | 0.0006 | 0.001 | 0.0014 | 0.0341 | 0.0449 | 0.0596 | 0.0368 | 0.0512 | 0.0427 | 0.0357 | 0.0496 | 0.0424 | 0.0447 | 0.0495 | 0.0473 |
| Glycolate | 0.009 | 0.0117 | 0.0109 | 0.0134 | 0.014 | 0.0153 | 0.0312 | 0.0397 | 0.0335 | 0.0297 | 0.044 | 0.0503 | 0.0079 | 0.0085 | 0.0096 | 0.0074 | 0.0106 | 0.0083 | 0.0087 | 0.0092 | 0.0066 |
| Guanidoacetate | 0.0163 | 0.0144 | 0.0161 | 0.0193 | 0.0204 | 0.0227 | 0.0281 | 0.0324 | 0.0296 | 0.0193 | 0.0257 | 0.0303 | 0.0233 | 0.0291 | 0.027 | 0.0225 | 0.0276 | 0.0212 | 0.0261 | 0.0488 | 0.0223 |
| Histamine | 0.0004 | 0.001 | 0.0021 | 0.012 | 0.0144 | 0.0172 | 0.0044 | 0.0049 | 0.0067 | 0.004 | 0.0052 | 0.0069 | 0.008 | 0.0122 | 0.0182 | 0.0251 | 0.0194 | 0.013 | 0.0052 | 0.0055 | 0.0064 |
| Isoleucine | 0.0014 | 0.0018 | 0.0023 | 0.0034 | 0.004 | 0.0049 | 0.0133 | 0.0189 | 0.0175 | 0.0085 | 0.0186 | 0.021 | 0.0427 | 0.056 | 0.049 | 0.0432 | 0.0518 | 0.0414 | 0.0496 | 0.0494 | 0.0593 |
| Lactate | 0.0108 | 0.0088 | 0.0074 | 0.0611 | 0.031 | 0.0308 | 0.1749 | 0.2272 | 0.2034 | 0.1205 | 0.1787 | 0.2067 | 0.1386 | 0.161 | 0.1539 | 0.1228 | 0.172 | 0.1549 | 0.1492 | 0.0907 | 0.1389 |
| Leucine | 0.0018 | 0.0021 | 0.0026 | 0.0017 | 0.0036 | 0.0051 | 0.0311 | 0.0393 | 0.0331 | 0.0141 | 0.021 | 0.0388 | 0.022 | 0.0388 | 0.0322 | 0.0201 | 0.0236 | 0.0131 | 0.0282 | 0.0372 | 0.0305 |
| Maleate | 0.0002 | 0.0003 | 0.0004 | 0.0003 | 0.0004 | 0.0007 | 0.0007 | 0.0008 | 0.0009 | 0.0008 | 0.001 | 0.0006 | 0.0012 | 0.0022 | 0.0024 | 0.0009 | 0.0017 | 0.0022 | 0.0025 | 0.001 | 0.0015 |
| Malonate | 0.0009 | 0.0013 | 0.0011 | 0.0019 | 0.0025 | 0.0039 | 0.0041 | 0.0034 | 0.0055 | 0.0036 | 0.0032 | 0.0041 | 0.0064 | 0.0095 | 0.0085 | 0.0077 | 0.0118 | 0.0113 | 0.0091 | 0.0089 | 0.0099 |
| Methionine | 0.0003 | 0.0003 | 0.0005 | 0.0004 | 0.0007 | 0.0013 | 0.0011 | 0.0013 | 0.002 | 0.0031 | 0.0041 | 0.0058 | 0.0026 | 0.0025 | 0.0034 | 0.0031 | 0.0041 | 0.0034 | 0.0041 | 0.0062 | 0.0048 |
| N-Acetylcysteine | 0.0012 | 0.0016 | 0.0029 | 0.0099 | 0.0081 | 0.0152 | 0.004 | 0.0038 | 0.0064 | 0.002 | 0.0027 | 0.0019 | 0.0029 | 0.0028 | 0.0022 | 0.0029 | 0.0038 | 0.0028 | 0.0046 | 0.0007 | 0.0038 |
| N-Acetylglucosamine | 0.0006 | 0.0008 | 0.0013 | 0.0018 | 0.0028 | 0.0035 | 0.0084 | 0.0091 | 0.0122 | 0.0045 | 0.0056 | 0.0068 | 0.0066 | 0.0069 | 0.0076 | 0.0068 | 0.0101 | 0.0063 | 0.0105 | 0.0088 | 0.0087 |
| N,N-Dimethylglycine | 0.0001 | 0.0007 | 0.0004 | 0.0005 | 0.0009 | 0.0016 | 0.0021 | 0.009 | 0.0028 | 0.0047 | 0.0056 | 0.0065 | 0.0059 | 0.0058 | 0.0065 | 0.006 | 0.0063 | 0.0063 | 0.0054 | 0.0061 | 0.0058 |
| NADH | 0.0034 | 0.0025 | 0.0016 | 0.0041 | 0.003 | 0.0037 | 0.0027 | 0.0032 | 0.0059 | 0.0016 | 0.0028 | 0.0025 | 0.0021 | 0.0025 | 0.0019 | 0.0031 | 0.0044 | 0.0032 | 0.0081 | 0.0105 | 0.009 |
| NADPH | 0.0179 | 0.0146 | 0.0128 | 0.0292 | 0.0315 | 0.0343 | 0.0223 | 0.0266 | 0.0287 | 0.0127 | 0.0211 | 0.0218 | 0.0152 | 0.0119 | 0.0124 | 0.0108 | 0.0181 | 0.0135 | 0.0149 | 0.0118 | 0.0158 |
| Propionate | 0.0211 | 0.0285 | 0.0289 | 0.0489 | 0.0508 | 0.0566 | 0.1123 | 0.1945 | 0.129 | 0.1005 | 0.0814 | 0.0909 | 0.0459 | 0.0931 | 0.0715 | 0.0455 | 0.0678 | 0.0525 | 0.0589 | 0.0687 | 0.0687 |
| Putrescine | 0.0007 | 0.0009 | 0.0018 | 0.0034 | 0.005 | 0.006 | 0.018 | 0.0106 | 0.0127 | 0.0179 | 0.0241 | 0.0275 | 0.0485 | 0.0458 | 0.0292 | 0.0389 | 0.0458 | 0.0363 | 0.0433 | 0.049 | 0.0607 |
| Pyruvate | 18.2397 | 13.4348 | 14.4514 | 9.3247 | 9.7475 | 10.3671 | 0.0028 | 0.0067 | 0.0066 | 0.0021 | 0.0027 | 0.0034 | 0.0023 | 0.0036 | 0.0038 | 0.0033 | 0.0046 | 0.0048 | 0.0045 | 0.0043 | 0.004 |
| Serine | 0.0032 | 0.0056 | 0.0079 | 0.0135 | 0.0115 | 0.0145 | 0.0208 | 0.0549 | 0.0501 | 0.0337 | 0.0414 | 0.0499 | 0.0468 | 0.0585 | 0.0578 | 0.0214 | 0.0196 | 0.0144 | 0.0435 | 0.0537 | 0.0379 |
| Succinate | 0.0002 | 0.0006 | 0.0013 | 0.0012 | 0.002 | 0.0034 | 0.0023 | 0.003 | 0.0034 | 0.0011 | 0.0016 | 0.0013 | 0.0016 | 0.0023 | 0.002 | 0.0018 | 0.0025 | 0.0022 | 0.0021 | 0.0026 | 0.0019 |
| Threonine | 0.0004 | 0.0004 | 0.0007 | 0.0058 | 0.0098 | 0.0067 | 0.0102 | 0.0121 | 0.0174 | 0.0184 | 0.0128 | 0.0223 | 0.0915 | 0.1051 | 0.1237 | 0.0339 | 0.0359 | 0.0258 | 0.0166 | 0.051 | 0.0151 |
| Thymine | 0.0001 | 0.0002 | 0.0003 | 0.0005 | 0.0005 | 0.0008 | 0.0008 | 0.0008 | 0.0011 | 0.0013 | 0.0021 | 0.0009 | 0.0026 | 0.0038 | 0.0028 | 0.0019 | 0.0025 | 0.002 | 0.0034 | 0.0044 | 0.0027 |
| Uracil | 0.0001 | 0.0002 | 0.0002 | 0.0005 | 0.0002 | 0.0009 | 0.0027 | 0.0006 | 0.0009 | 0.0017 | 0.0026 | 0.0019 | 0.0007 | 0.0011 | 0.0014 | 0.0009 | 0.0008 | 0.0013 | 0.0039 | 0.0049 | 0.0047 |
| Valine | 0.051 | 0.0332 | 0.0369 | 0.0742 | 0.0746 | 0.0909 | 0.9529 | 1.2929 | 1.1076 | 0.6338 | 0.9429 | 1.2047 | 0.7109 | 0.9199 | 0.8546 | 0.644 | 0.9079 | 0.91 | 0.5389 | 0.4534 | 0.5223 |

Additionally, several metabolites related to amino acid biosynthesis and protein production, such as isoleucine, valine, and citrate, form another cluster. The clustering of these metabolites suggests that they are closely linked to the metabolic demands of recombinant protein synthesis, especially following IPTG induction when the demand for amino acids sharply increases. This cluster also includes succinyl-CoA, an intermediate in the TCA cycle, further indicating the importance of energy production and carbon flow through central metabolism to support protein biosynthesis.

The hierarchical clustering dendrogram (Figure 4B) further elucidates the relationships between metabolites based on their correlation profiles. The dendrogram groups metabolites into clusters according to their similarity in correlation patterns, with the vertical axis representing the distance (or dissimilarity) between clusters. Metabolites with similar roles or functions are grouped together, indicating shared regulatory mechanisms or metabolic pathways. At the top of the dendrogram, pyruvate and acetate form a distinct cluster, reflecting their central roles in energy metabolism and overflow metabolism, respectively. The strong correlation between pyruvate and acetate likely reflects the bacterial metabolism’s response to the high flux of carbon through glycolysis and the citric acid cycle, leading to acetate secretion as a byproduct during later stages of growth. Interestingly, metabolites involved in nucleotide metabolism, such as adenine and uracil, cluster together, reflecting their role in DNA and RNA synthesis, which may be essential for supporting bacterial growth and protein production during the exponential phase. This cluster also includes metabolites like thymine, emphasizing the coordinated regulation of nucleotide metabolism in response to the energy and biosynthetic demands imposed by recombinant protein production. Overall, the hierarchical clustering reveals that metabolite profiles are organized into functional groups based on their roles in central carbon metabolism, amino acid biosynthesis, and energy production. The tight clustering of key metabolites involved in pyruvate metabolism and BCAA biosynthesis highlights the importance of these pathways in supporting bacterial growth and protein production under minimal medium conditions with pyruvate as the sole carbon source. The Pearson correlation heatmap and hierarchical clustering dendrogram together reveal distinct metabolic modules that are co-regulated during bacterial growth and recombinant protein production. Metabolites involved in energy metabolism, amino acid biosynthesis, and nucleotide metabolism show strong correlations, reflecting the bacterial cells’ need to balance energy production with biosynthesis to support protein production. These findings provide a comprehensive overview of how central metabolism is reprogrammed in response to the metabolic demands imposed by recombinant protein synthesis.

## Discussion and conclusions

The metabolic analysis provides significant insights into the adaptations of *E. coli* when grown with pyruvate as the sole carbon source. Our results highlight the complex balance between central carbon metabolism and overflow pathways under both basal conditions and recombinant protein production.

### Pyruvate Utilization and Overflow Metabolism

Pyruvate, as expected, was rapidly consumed by the bacterial cells, with its concentration significantly reduced well before IPTG induction. This early depletion of pyruvate suggests its immediate utilization in key metabolic pathways such as glycolysis, the citric acid (TCA) cycle, and amino acid biosynthesis. The pivotal role of pyruvate in *E. coli* metabolism is well established, serving not only as an energy source but also as a precursor for various biosynthetic reactions that are critical for cell growth and maintenance (Zhang et al., 2019). Our data are consistent with previous studies that have demonstrated the versatile role of pyruvate in driving both energy production and biosynthesis during the exponential phase of bacterial growth (Zhu et al., 2020).

Prior to IPTG induction, we observed significant accumulation of both acetate and lactate in the culture medium, indicating that excess pyruvate was being diverted into overflow metabolic pathways. This phenomenon, where surplus pyruvate is converted to byproducts such as acetate and lactate, is well documented in rapidly growing bacterial cultures, particularly when carbon availability exceeds the oxidative capacity of the TCA cycle (Holms, 1986). The accumulation of these metabolites in the extracellular environment suggests that *E. coli* cells are expelling them to prevent intracellular metabolic imbalances. Once expelled, the re-uptake and subsequent metabolism of acetate and lactate appear to be inefficient, which could explain why their concentrations remain elevated throughout the experiment. This observation is in line with studies reporting that the re-assimilation of acetate and lactate by *E. coli* is energetically costly and tightly regulated (Vemuri et al., 2006; Wolfe, 2005). The inability of the cells to efficiently reclaim acetate and lactate likely reflects a strategy to maintain metabolic homeostasis by eliminating surplus carbon. Once pyruvate is converted to these byproducts and secreted, their re-utilization may be energetically unfavorable, particularly under conditions where the TCA cycle is already operating at full capacity. This highlights the overflow metabolism phenomenon, which is exacerbated when pyruvate availability exceeds the oxidative needs of the cells (Basan et al., 2015). The persistent high levels of acetate and lactate in the medium further suggest that the cells prioritize maintaining intracellular balance over attempting to reclaim and metabolize these byproducts, thus allowing their excretion into the extracellular environment (Enjalbert et al., 2015).

### Metabolic Shifts Post-IPTG Induction

Following IPTG induction at 9 hours, we observed that the concentrations of acetate and lactate remained elevated. This continued presence of overflow metabolites suggests that the metabolic burden imposed by recombinant protein production does not alleviate overflow metabolism, but rather intensifies it. During high metabolic demands, such as during recombinant protein synthesis, the increased carbon flux through glycolysis and the TCA cycle may further limit the cells’ ability to fully oxidize pyruvate, leading to continued diversion into acetate and lactate pathways (El-Mansi et al., 2006). Citrate, a key intermediate of the TCA cycle, is essential for energy generation and the production of biosynthetic precursors. Its accumulation suggests that the cells are attempting to balance energy production with the increased demand for biosynthetic precursors necessary for protein synthesis (Luo et al., 2020). This finding is consistent with earlier work demonstrating that, under recombinant protein production conditions, bacterial cells often increase TCA cycle activity to meet the elevated demand for ATP and precursor metabolites (Gill et al., 2000). Despite this increased TCA cycle flux, the persistent high levels of acetate and lactate indicate that the cells are unable to fully process the available pyruvate through oxidative pathways, leading to its continuous conversion into overflow metabolites. This metabolic response likely represents a physiological adaptation to the stress imposed by recombinant protein production, where the diversion of pyruvate into overflow pathways helps to prevent metabolic overload within the cell (Basan et al., 2015).

### Implications for Bioprocess Optimization

The results of this study have several important implications for optimizing bacterial growth conditions in biotechnological applications. The accumulation of acetate and lactate can have deleterious effects on bacterial growth and protein production, as these byproducts contribute to acidification of the culture medium and inhibit key cellular processes (Enjalbert et al., 2015). Strategies to mitigate the accumulation of these byproducts, such as optimizing pyruvate feeding rates or engineering strains with improved acetate and lactate assimilation capabilities, could enhance the efficiency of recombinant protein production (Vemuri et al., 2006; Sauer et al., 2011). Additionally, the findings underscore the importance of fine-tuning carbon fluxes in metabolic pathways to minimize the diversion of carbon into overflow metabolites. Engineering bacterial strains with an enhanced capacity for pyruvate oxidation, or increasing the TCA cycle’s capacity to process higher fluxes of carbon, could help reduce the reliance on overflow pathways, thereby improving growth and protein production yields (Holms, 1986). Further investigation into strain engineering strategies that enhance carbon utilization while reducing byproduct formation could prove beneficial for large-scale industrial fermentation processes (Sauer et al., 2011).

This study provides critical insights into the metabolic behavior of *E. coli* during recombinant protein production when grown on pyruvate as the sole carbon source. The rapid depletion of pyruvate, coupled with the significant accumulation of acetate and lactate in the medium, underscores the challenges associated with pyruvate over-supplementation. Excess pyruvate, while serving as a critical substrate for energy production and biosynthesis, is inefficiently utilized under these conditions, leading to its diversion into overflow pathways and the excretion of byproducts. These findings highlight the importance of fine-tuning carbon source supplementation in biotechnological processes, as supplying pyruvate in excessive amounts can lead to metabolic imbalances and the inefficient use of carbon resources. By optimizing the levels of pyruvate provided and improving the metabolic capacity of the cells to process it, the accumulation of waste byproducts like acetate and lactate can be minimized, ultimately enhancing the efficiency of recombinant protein production. Future work should focus on strain engineering and process optimization to better manage carbon fluxes, reduce waste, and improve the overall yield and productivity of industrial fermentation systems.

## ACKNOWLEDGEMENT

The authors acknowledge the use of the services and facilities of n^2^STAR-Koç University Nanofabrication and Nanocharacterization Center for Scientific and Technological Advanced Research. KK acknowledges support from TU BI TAK (2209-A-December 2021).

## Author contributions

Ç.D.: Investigation, Conceptualization, Methodology, Supervision, Writing-Original draft preparation, Formal analysis, Writing−Reviewing and Editing K.K.:Investigation

## Data availability

The datasets generated during and/or analyzed during the current study are available from the corresponding author on reasonable request.

## Declarations

The authors declare no competing interests.

